# Attachment performance of cuttlefish (*Sepia officinalis*) suckers depends on the interaction between papillae and substrate topography

**DOI:** 10.1101/2025.08.05.668641

**Authors:** Jan Severin te Lindert, Brett Klaassen van Oorschot, Tabo Geelen, Esther L. te Lindert Blommert, Marcel Giesbers, Baowen Zhang, Yoerick T. Lankhof, Sander W.S. Gussekloo, Florian T. Muijres, Guillermo J. Amador

**Author notes:** Co-correspondence. Co-first authorship (Jan Severin te Lindert, Brett Klaassen van Oorschot).

## Abstract

Cephalopods are versatile predators, with many species using suckers to capture prey. These suckers attach to substrates ranging from the stiff, rough exoskeletons of crustaceans to the soft, smooth tissues of other cephalopods. Despite generating higher suction pressures than octopuses, less attention has been given to the biomechanics of cuttlefish suckers. Cuttlefish suckers exhibit a stiff, rough papillated rim that acts as a seal. We hypothesise that these papillae have evolved to attach to rough substrates matching their rugosity to expel water at the contact interface. To test this hypothesis, we investigated the passive attachment performance of common cuttlefish (*Sepia officinalis*) suckers *ex vivo* on a variety of artificial substrates that differed in stiffness and roughness. We found that sucker attachment forces varied significantly with substrate roughness, with the highest forces occurring on roughnesses that coincided with the average sucker papillae height (root-mean-square ∼ 6.84 µm). Additionally, attachment forces were higher on stiffer substrates. We developed a mathematical model for predicting leakage across the sucker rim that qualitatively agrees with our experimental results. These findings indicate that the papillae may aid attachment performance, particularly to natural prey, and could inform the design and development of versatile bioinspired suction cups.

## 1 Introduction

Cephalopods, like the common cuttlefish (*Sepia officinalis*), are highly versatile predators that use their sucker-lined tentacles and arms to capture and manipulate prey, such as fish, crabs, shrimps and other cephalopods (Fig. 1; [1–5]). The skin of these prey (i.e., exoskeletons, scales, and soft tissues) vary both in stiffness (how much an object resists deformation from an external force) and roughness (the deviation in the normal vector of a surface) [6–8]. Hence, effective attachment to a variety of substrates is crucial for survival. Rough substrates present challenges because of increased risk of leakage at the sucker rim, and soft substrates can trap pockets of fluid at the contact interface that hinder adhesion and friction [9, 10]. Therefore, the sucker rim, through which contact and sealing of the sucker cavity are mediated, is chiefly responsible for the stability of suction during attachment.

**Figure 1:**
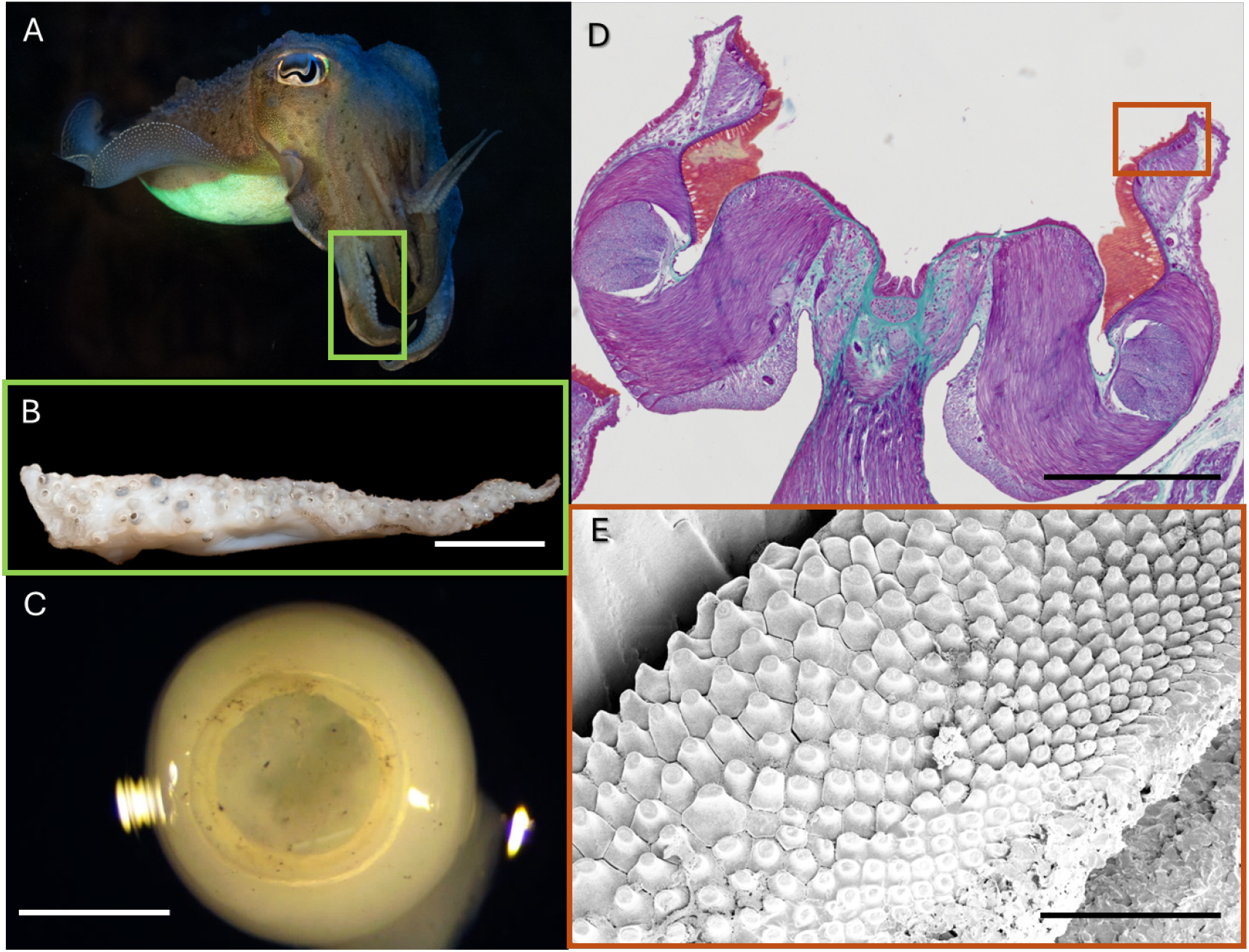
(A) Common cuttlefish (*Sepia officinalis*); (B) Excised arm with suckers (scale bar represents 2 cm); (C) Close-up of a sucker (scale bar represents 1 mm); (D) Histology image of a sucker (scale bar represents 500 µm), with papillae surface highlighted with a red box; and (E) Scanning electron micrograph (SEM) of the papillae on the sucker rim (scale bar represents 100 µm).

To attach successfully, suckers need to conform to rough and irregular features to provide the seal needed for suction. For example, in clingfish (Gobiesocidae), relatively soft suckers have evolved with microstructures (papillae) that conform around the features of the substrate, providing a tight seal that enables them to support up to 230 times their body weight [11, 12]. Depending on the specific morphology of these specialised papillae, they are hypothesised to enhance friction or mechanical interlocking, or both [13]. However, the soft suckers of northern clingfish (*Gobiesox maeandricus*) struggle to adhere to soft substrates [14]. It has been suggested that the combined stiffness of two soft objects in contact (which, together are termed “effective modulus”) mediates attachment performance, where attachment performance increases with increasing effective modulus [14–16]. However, for suckers, a physical understanding of this mechanism is missing, especially when large suction pressures are present.

As seen from natural hunting behaviour, the distinct morphology of cuttlefish suckers allows for secure attachment to both soft and rough substrates [1, 17, 18]. The suckers of cuttlefish exhibit three unique structures: 1) the stalk (or peduncle) that connects the sucker to the arm or tentacle; 2) the infundibular ring (also called “sucker teeth ring”), a stiff structure sometimes bearing sharp teeth; and 3) the papillated rim connected to the infundibular ring, which exhibits micrometre-scale papillae (Fig. 1D,E).

The cuttlefish sucker is hypothesised to use the stalk as a piston [19]. First, pushing the stalk towards the substrate and into the sucker expels water from the sucker cavity. Then, retracting the stalk generates suction by increasing cavity volume and decreasing pressure within the cavity. To avoid collapse of the sucker, it is hypothesised that the infundibular ring provides rigidity. Together, this allows for very strong attachment, up to tenacities (adhesive pressure as force per unit contact area, Eqn. 1) of 800 kPa, as a previous study has shown for smooth, stiff substrates (glass) in water [19]. However, the suckers’ attachment performance on soft and rough substrates and the role of the papillae have yet to be investigated.

The morphology of the cuttlefish sucker provides an interesting avenue for the development of bioinspired suction cups. Various other bioinspired suction cups based on, for example, octopus, clingfish, and remora suckers have demonstrated secure attachment on various substrates [20–24]. However, cuttlefish suckers have not yet been examined as a source of bioinspiration for synthetic suction cups, which is surprising given their remarkable attachment performance [19].

Here, we quantified the attachment performance of *Sepia officinalis* arm suckers on a range of artificial substrates differing in stiffness and roughness. We defined three complementary hypotheses about the functional role of papillae on suction performance: First, similar to the microstructures exhibited by clingfish, we hypothesised that the papillae of cuttlefish suckers aid in attachment to rough substrates by conforming to substrate roughness. Secondly, we hypothesised that the stiffness of the papillae may enable secure attachment onto soft substrates, as a stiffer rim reduces the likelihood of water entrapment at the rim interface [9]. Finally, we hypothesised that cuttlefish suckers achieve highest attachment forces on stiff substrates whose roughness matches that of the papillae.

We tested these three hypotheses using a combined experimental and modelling approach. We quantified sucker attachment performance using a series of attachment experiments with excised suckers from the arms of *S. officinalis*. For these experiments, we used artificial silicone rubber substrates with systematically varying stiffnesses and roughnesses, as was similarly done previously with clingfish [14]. We then measured and compared the roughnesses of the papillae to the artificial substrates and natural occurring prey species: crab (*Carcinus maenas*), shrimp (*Crangon crangon*) and fish (*Solea solea*), as well as stiffness values from the literature. Finally, we used a mathematical model of leakage through the papillated rim to understand how the interaction between papillae and substrate causes the measured variations in sucker attachment performance.

## 2 Materials and Methods

### 2.1 Animals

In this study, suckers of 19 European common cuttlefish (*Sepia officinalis*) were used for attachment performance measurements (pull-off force experiment, described later in this section). Cuttlefish and prey species, shrimp (*Crangon crangon*) and crab (*Carcinus maenas*), were obtained from fisheries (Visafslag Scheveningen B.V.) in Scheveningen, The Netherlands. The samples were caught and kept on ice for ∼24 hours, then weighed and kept frozen in airtight bags at −20°C until the start of the experiment, limiting the specimen to one freeze-thaw cycle. The duration that specimens were kept frozen did not influence adhesive performance (see Fig. S1A). Freshly deceased sole (*Solea solea*) were obtained from the Wageningen University animal facility in Wageningen, The Netherlands, and stored in a similar manner.

Dead tissue was used because we were interested in passive properties of the suckers and live or freshly killed animals can still have unwanted reflexive muscle activity. Moreover, previous research has shown that freezing has limited effects on the biomechanical properties of passive, connective tissues [25–28]. Consistent with this, our tests demonstrate that cuttlefish sucker attachment performance on glass is comparable between *in vivo* [19], frozen squid suckers [28], and our own samples (see Fig. S1B).

### 2.2 Sucker attachment experiments

To quantify the attachment performance of cuttlefish arm suckers, *ex vivo* experiments were performed with single excised suckers from the arms of *S. officinalis*. The suckers had an average outer diameter of 3.6 mm ± 0.6 mm (mean ± s.d.). These suckers were tested on artificial substrates with various stiffnesses and roughnesses. Statistical modelling was used to correlate the variations in adhesive performance with the variations in substrate characteristics.

#### 2.2.1 Producing and characterizing the artificial substrates

In total, 12 different substrates were made, varying in both roughness (4 types) and stiffness (3 types). Three silicone rubbers with different stiffnesses were used: Ecoflex^™^ 00-30 (“Soft”), Dragon Skin^™^ 30 (“Medium”), and Sylgard^™^ 184 (“Stiff”), which have elastic moduli *E* of 70 kPa, 590 kPa, and 1320 kPa, respectively, as reported by the manufacturers (Smooth-On Inc., DOW Sylgard^™^) [29]. Silicones were mixed following the manufacturers’ instructions and degassed in a vacuum chamber until trapped air bubbles were removed. 3D-printed moulds were based on a 12-mm diameter borosilicate lens with a radius of curvature of 37.22 mm and thickness of 2 mm from Edmund Optics® [30]. To create different roughness values, smooth lapping film or one of three types of 3M® sandpaper were applied to the bottom of the mould: P1200, P240, and P80, corresponding to a roughness (*Sq*, root mean square) of *Sq* = 0.08 *µ*m (“Smooth”), 5.35 *µ*m (“Fine”), 13.27 *µ*m (“Semi-coarse”), and 60.32 *µ*m (“Coarse”). The different silicones were cast directly into these moulds and cured at room temperature (18°C) for 24–72 hours, following manufacturer guidelines. These substrates were glued (3M® Scotch-Weld^™^ cyanoacrylate) onto a 3D-printed puck (5 mm height and 12 mm diameter) made of polylactic acid (PLA), acting as a stiff backing layer (Fig. 2A).

**Figure 2:**
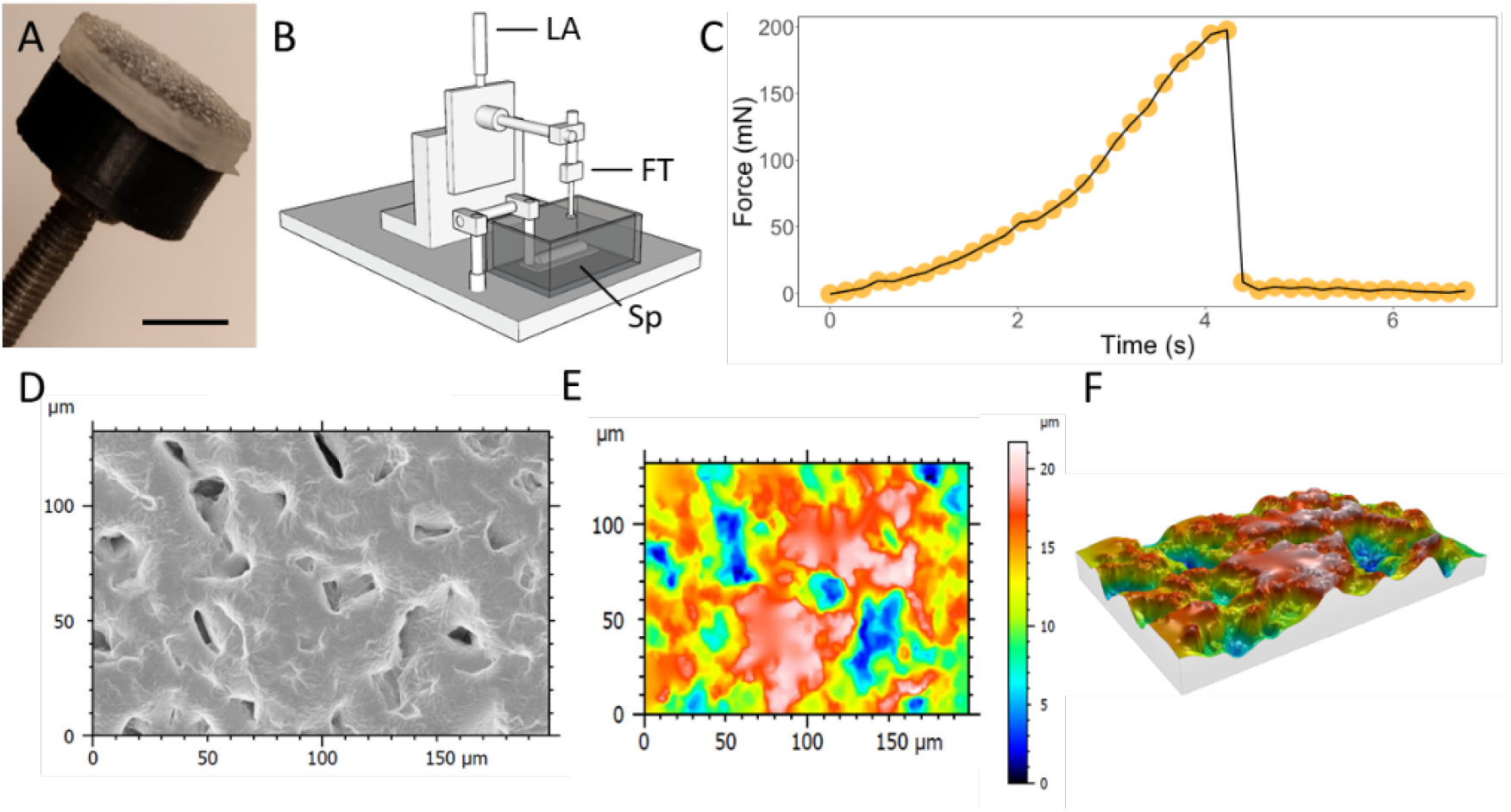
(A) Image of the artificial substrate connected to the 3D-printed puck used for attachment experiments. Scale bar represents 5 mm; (B) Schematic of the experimental setup for measuring attachment force. LA: linear actuator, FT: force transducer, and Sp: specimen; (C) Typical plot from an attachment force measurement; (D) Scanning electron micrograph (SEM) images of the coarse substrate; (E) Contour plot of roughness distribution of the coarse substrate from D; (F) Topology of the coarse substrate from D; E-F were obtained using stereoscopic SEM scans and are color-coded with substrate height (see color bar in E).

#### 2.2.2 Attachment experiments

The attachment experiments were performed using a custom-designed experimental setup (Fig. 2B). It consists of a container for artificial seawater with a cuttlefish arm holder and a linear stage. The linear stage includes a linear actuator (Thorlabs® Z825B) for moving the substrate pucks and a submersible force transducer (FUTEK® LSB210 100-gram capacity) for measuring the sucker attachment forces. To reduce external measurement noise, the whole setup was placed on a vibration isolation table.

Prior to each experiment, one of the twelve substrates was mounted onto the linear stage and a defrosted cuttlefish arm was placed in the plastic transparent container filled with 35 g L^-1^ seawater solution (Instant Ocean®) and ice to keep it cool. The cuttlefish arm was cut to flatten it and was then attached on the frosted side of a Superfrost^™^ Microscope Slide (Epredia®) using cyanoacrylate glue (3M® Scotch-Weld^™^).

Upon the start of the experiment, the mounted substrate was lowered at 1 mm s^-1^ using the motorized linear actuator until contact was achieved between substrate and sucker, and a compressive preload force of 200 mN was applied. This preload was chosen based on preliminary data from cuttlefish prey strikes, which showed that the preload applied during tentacle strikes is in the range of 100–200 mN. Upon reaching the preload force, the indenter stopped for 2 seconds, after which the actuator retracted at 2 mm s^-1^ until the sucker detached from the substrate. This procedure is consistent with previous measurements on octopus suckers [30]. The use of substrate samples was pseudo-randomized and, since not all suckers stuck, the test was repeated until a suction event was measured. To estimate the necessary sample size, an *a priori* power analysis was performed using Cohen’s *f* estimated from a generalised linear model (GLM) fitted to pilot data, including main effects and interactions of stiffness and roughness.

From each successful trial, the peak force *F* just before detachment was extracted. A typical plot of the force measurements during detachment is shown in Fig. 2C. Based on the peak adhesive force, we then calculated the tenacity *P* (i.e., adhesive pull-off pressure) as

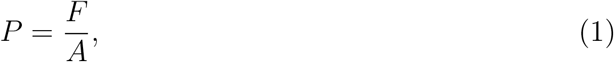

where *F* is the peak adhesive force and *A* is the inner area of the sucker’s infundibular ring, as was done in previous experiments [14, 19]. The contact area for each tested sucker was measured after the experiment using a dissection scope and camera (Amscope® MU1403B) and imageJ 1.53t [31].

### 2.3 Histology

Histology was used to visualise the distinct morphology of a cuttlefish sucker. A modified Masson’s trichrome stain was used to stain musculature, connective tissue, and proteinaceous tissue (Kiernan, 2015). Cuttlefish arms, excised from thawed cuttlefish, were fixed in a 4% formalin solution in phosphate-buffered saline (PBS). After embedding in paraffin, 6 ™ 10 *µ*m sections were cut with a Leica® Jung Biocut RM2035 microtome and put onto Superfrost™ Plus Adhesion microscope slides (Epredia®). Slides were deparaffinized twice in xylene for 5 minutes and then rehydrated using an ethanol series of 100%, 100%, 96%, 90%, 80% and 70% for 3 minutes each, followed by 2x demineralised water for 5 minutes each. Slides were then stained with a modified Masson’s trichrome staining: submerged in Mayer’s haematoxylin solution for 5 minutes, rinsed under running tap water for 10 minutes, dipped in acid fuchsin and Orange G for 3 seconds, submerged in demineralised water for 2 minutes, submerged in phosphotungstic acid (5%) for 4 minutes, demineralised water for 2 minutes, Light Green SF (1%) for 10 minutes, and two times in demineralised water for 1 minute. After staining, we washed 4 times for 2 minutes in 100% ethanol. Slides were then washed 3 times for 2 minutes in xylene and then covered with a cover slip and sealed with DEPEX mounting media. Samples were imaged using a Leica® DM6b microscope with a DFC450c camera and the LASX program. The histological slide is shown in (Fig. 1D).

### 2.4 Scanning electron microscopy (SEM)

Roughness of papillae, substrates, and prey was quantified using scanning electron microscopy (SEM). In preparation for SEM, thawed cuttlefish suckers were suspended in fixative (2.5% glutaraldehyde and 2% paraformaldehyde in 0.1M phosphate buffer) to crosslink proteins and then left overnight at room temperature. The suckers were washed 6 times for 15 minutes in 0.1M phosphate buffer. Subsequently, the suckers were put in 1% osmium tetroxide in 0.1M phosphate buffer and left for 1 hour at room temperature to crosslink fats. The specimens were washed 3 times with ultra-purified water and dehydrated through fixation in an ascending ethanol series (10%, 30%, 50%, 70%, 80%, 90%, 96%, 100%) for 15 minutes each and the last step was repeated three times. Then, samples were critical-point dried (Leica® CPD 300, Vienna, Austria) followed by sputter-coating with a 12 nm tungsten layer at three angles (−45 degrees, 0, and 45 degrees; Leica® SCD 500, Vienna, Austria) to ensure good conductivity. The same preparations were used for the substrate of the different prey species: shrimp carapace and first abdominal segment (*Crangon crangon*), fish dorsal side (*Solea solea*) and crab both ventral and dorsal carapace (*Carcinus maenas*). However, the osmium tetroxide fixation step was omitted as the amount of fat tissue is negligible for the fish scales and exoskeletons of these species.

The four stiff silicone rubber substrates varying in roughness (coarse, semi-coarse, fine, and smooth) were mounted onto aluminium SEM stubs using a very thin layer of carbon adhesive glue (Leit-C, EMS, Hatfield, PA, USA). The samples were kept at near-zero vacuum pressure overnight to dry the Leit-C glue. Subsequently, the substrates were sputter-coated with 12 nm of tungsten at two angles (+45° and −45°) to ensure good conductivity (Leica® SCD 500, Vienna, Austria).

Field-emission SEM (FEI® Magellan 400, Eindhoven, The Netherlands) was used to validate and measure the roughness of the substrates, prey species, and the papillae size of the cuttlefish suckers. Images were captured with the secondary electron detector set at 2 kV and 13 pA. Images were taken at +3° and −3°, allowing for the generation of 3D stereoscopic images and subsequent measurement of the substrate topology (see Fig. 2D-F and Fig. S2). The images were obtained at 250× magnification for coarse stiff, semi-coarse stiff and fine stiff substrates. For the papillae and smooth stiff substrate, magnifications of 2500× and 5000×, respectively, were used to ensure adequate resolution. Similarly, prey scans were obtained at the following magnifications: sole at 250× and 2500×, crab at 2500×, and shrimp at 2500×. Characteristic SEM images of the substrates, prey specimens, and papillae are shown in Fig. 3.

**Figure 3:**
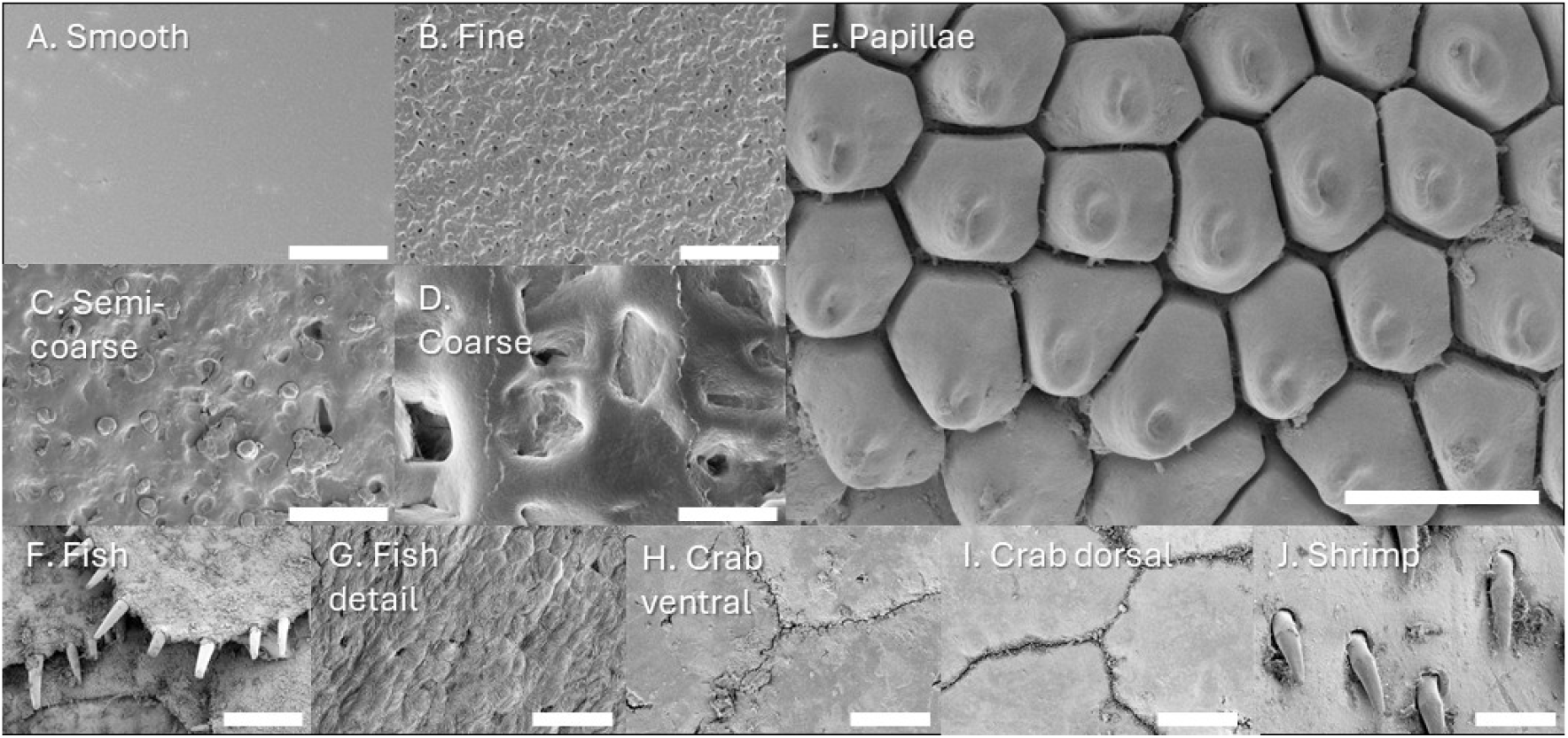
Scanning electron micrograph (SEM) images of: (A-D) the artificial substrates, (E) cuttlefish sucker papillae, and (F-J) the main prey groups. These prey include (F, G) fish (*Solea solea*), (H, I) crab (*Carcinus maenas*), and (J) shrimp (*Crangon crangon*). Scale bars represent a length of (A-D, F) 300 µm, (E, G-J) 30 µm.

### 2.5 Cryogenic Scanning Electron Microscopy

Cryogenic scanning electron microscopy (cryo-SEM) was used to visualise the interface between the papillae and fine substrate. The fine substrate was mounted on a holder using a thin layer of Tissue-Tek (EMS, Washington, PA, USA), after which a cuttlefish sucker was clamped on top of the fine substrate. The opening of the holder was slowly narrowed to press the sucker onto the substrate without deforming the sucker ring. As the silicone rubber fine substrate could not be broken with cryo-breaking, a cross-section was made at room temperature by cutting the sucker and substrate with a razor blade. After being plunged into liquid nitrogen, the sample was transferred to a cryo-preparation chamber (MED 020/VCT 100, Leica®, Vienna, Austria) and sublimated for 3 minutes. The samples were then sputter-coated with an 8-nm layer of tungsten from three angles and transferred under vacuum to the field emission scanning electron microscope (FEI® Magellan 400, Eindhoven, The Netherlands), where they were placed on the sample stage at –120°C. Images were captured using the secondary electron detector at 2 kV and 13 pA.

### 2.6 Analysis in MountainsSEM

Stereoscopic 3D reconstruction of the substrates was performed using the MountainsSEM® software (Digital Surf®, Besançon, France). During post-processing, outliers were removed using a proprietary software filter. Then, a 5-pixel smoothing filter was applied, and the image was levelled by fitting a least squares plane, which served as the reference plane. The root-mean-square height *Sq*, which corresponds to the standard deviation of the height distribution, was extracted from the height map with the following formula:

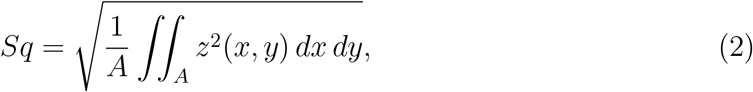

where *A* is the area, *x* and *y* are the plane coordinates, and *z* is the height of the substrate.

### 2.7 Statistical analyses

All statistical analyses were performed with R version 4.4.2 [32], and statistical results were reported as mean ± standard error of the model estimate, and the corresponding *p*-value. Effects and differences were assumed significant when *p* < 0.05.

#### 2.7.1 Tenacity analysis

Tenacity data were checked for normality using a Shapiro-Wilk test and QQ plots. To improve normality, a square root transformation was applied to the tenacity data prior to statistical analysis. Then, a linear mixed effects model (LMEM) with stiffness, roughness, and their interaction as main effects; sucker area as a fixed effect; and specimen ID as a random effect was used to examine their impact on tenacity. Sucker area was included because previous research has shown that smaller suckers produce significantly higher tenacities than larger suckers [19] (Fig. S3–S4). The interactions between stiffness and roughness were summarized with the emmeans R package [33]. Tenacity results were reported as the mean ± standard error of the LMEM estimates, with corresponding *p*-values.

#### 2.7.2 Roughness analysis

Differences in roughness between the artificial substrates, prey items and the cuttlefish papillae were examined as well. Due to unequal variances, a Welch’s ANOVA and a Games-Howell nonparametric post-hoc test was used to determine whether the substrates and prey items were significantly different in roughness from the cuttlefish papillae. Results on roughness effects were reported as the mean ± standard error with corresponding *p*-values.

### 2.8 Modelling of substrate-papillae interactions

#### 2.8.1 Mathematical model

While previous researchers have applied classical contact models of soft materials, or the Johnson-Kendall-Roberts (JKR) contact model, to gain insights into sucker rim-substrate interactions [14–16], these approaches only consider contact formation and neglect deformations caused by the significant suction pressures. Therefore, the direct implications of substrate stiffness on peak attachment performance are not fully captured with such models. We hypothesised that leakage across the papillated rim determined the maximum attachment performance of a sucker because maintaining a pressure difference between the inside and outside of the sucker cavity is essential for attachment. To test this hypothesis, we compared our results with a model in which the leakage (i.e., flow) between the papillae and substrate is simplified as pressure-driven plane Poiseuille flow between two infinite, parallel plates, as has been done previously for synthetic suction cups [34].

The papillae and substrate are represented as sine waves with the amplitude *a* and wavelength *λ* set equal to the roughness *Sq* measured for each, with the height *h* of each as a function of radial distance from the inside of the rim *x* being expressed as

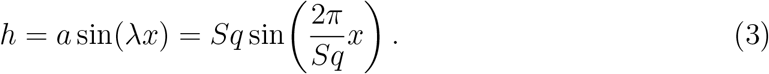

Since the substrate stiffness varies, we included substrate deformation due to the suction pressures generated by the suckers. Within the sucker, low pressures cause the substrate to deform. As a simplification, deformations to the asperities of the substrate have been neglected. This simplification may overestimate the leakage rate through softer substrates when papillae and substrate asperities are misaligned. The amount the substrate deforms was modelled assuming the substrate stretches like a spring, with the elastic modulus *E* as the spring constant. When the substrate is deformed, the part of the substrate in contact with the papillae is assumed to tilt by angle *θ*. In order to determine this tilting angle, the deformation of the substrate is assumed to be linear, where the substrate is pulled at the centre of the sucker and then sloped downwards towards the undeformed state outside of the sucker. Therefore, the angle *θ* can be expressed as

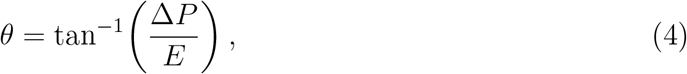

where Δ*P* = *P*_*o*_ ™ *P*_*i*_ is the pressure difference across the sucker rim with pressure outside *P*_*o*_ and inside *P*_*i*_ of the sucker. Δ*P* was assumed to be 100 kPa, which is the maximum practical pressure difference achievable without cavitation at sea level. While the pressure difference Δ*P* does vary depending on attachment performance, this simplification is made in order to make the model tractable. Therefore, the deformations may be slightly underestimated or overestimated, depending on the true attachment performance. A schematic of the mathematical model is shown in Fig. 4A.

**Figure 4:**
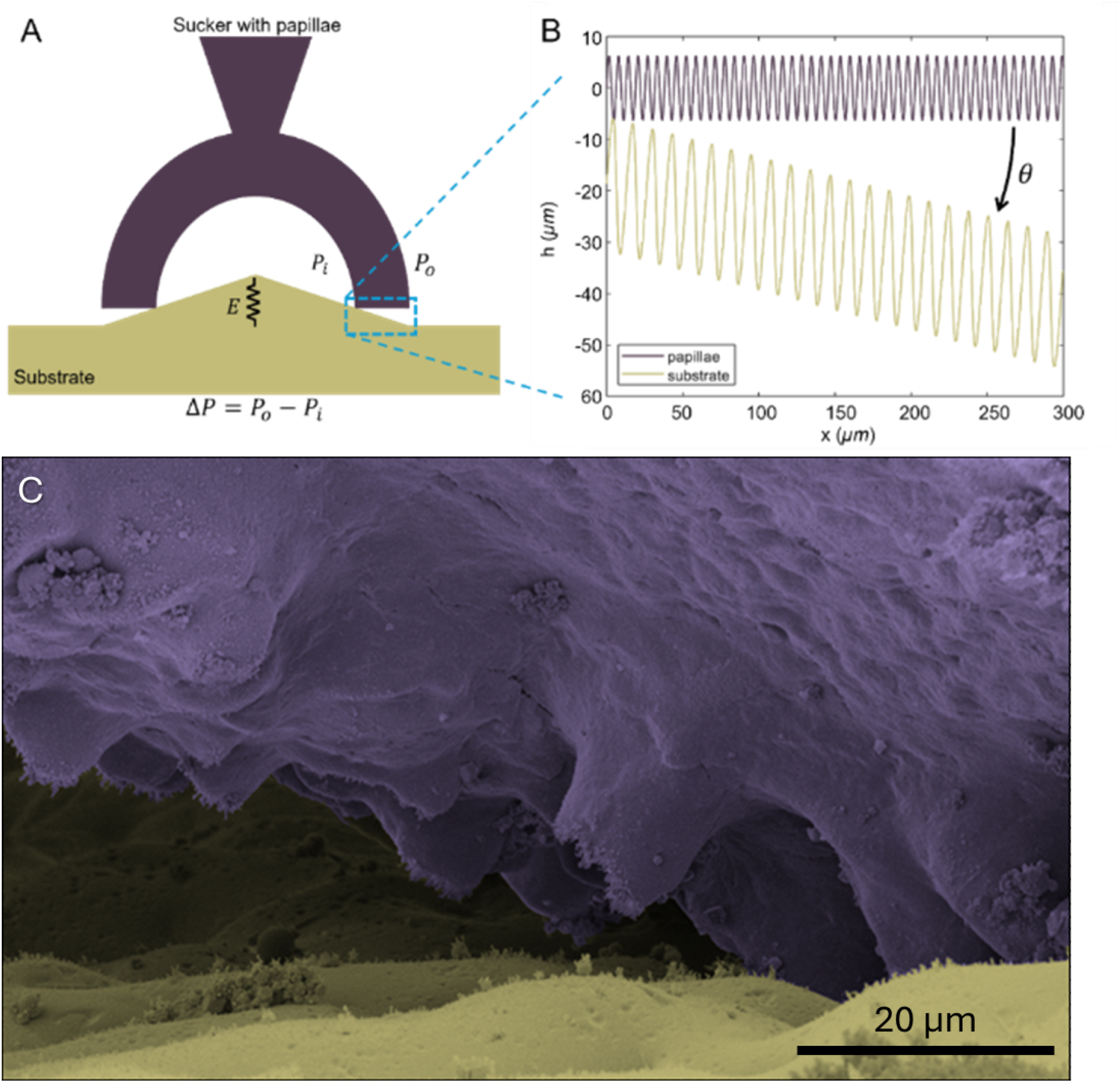
(A) Attachment model of the sucker with papillae to a rough compliant substrate, where *P*_*i*_ is the inner pressure, *P*_*o*_ is the ambient pressure, *E* is elastic modulus and spring constant of the substrate. (B) Plot of papillae and substrate modelled as sine waves, with *θ* being the tilt angle between the papillae and substrate due to deformation of the substrate caused by suction. (C) Visualisation of the interface between the papillae (purple) and the fine substrate (yellow) using cryo-SEM.

The contact of the papillae and substrate was represented by the two respective sine curves touching without overlap. An example of the sine curves representing the papillae and rough, deformed substrate is shown in Fig. 4B. Then, the average separation gap between the curves was calculated by subtracting them and taking an average. Finally, this separation gap *d* was used to determine how leakage flow rate through the sucker rim changes with substrate stiffness and roughness following the expression for plane Poiseuille flow. The flow rate of a fluid with viscosity *µ* between two parallel plates driven by a pressure gradient Δ*P/L*, where *L* = 300 *µ*m is the length across the sucker rim, is given as

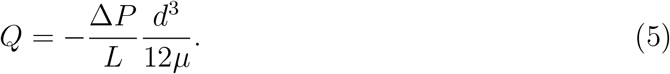

The alignment between the papillae and substrate asperities is stochastic, so the separation gap *d* was averaged for all possible alignments of the two representative curves.

#### 2.8.2 Finite element model

Finite element simulations were performed in Abaqus 2026 [35] to quantify the deformation of the substrate within the sucker under a suction pressure. To reduce computational cost, a static axisymmetric model was used. The substrate was modelled as a linearly elastic material with various stiffnesses *E* = 200 ™ 1400 kPa in increments of 200 kPa, while the Poisson’s ratio was fixed at 0.49.

The substrate was defined with a thickness of 2 mm and a radial extent of 4 mm. To represent the rigid contact boundary associated with the sucker ring, a rigid square with a side length of 0.5 mm was introduced, with its left edge located at a radial position of 1 mm. To improve numerical convergence, the lower-left corner of the rigid square was rounded with a fillet radius of 0.1 mm.

To reproduce the experimental constraints, the bottom surface and outer side surface of the substrate were fixed, corresponding to the glued boundaries in the experiments. The rigid square was fully constrained. A uniform outward gauge pressure of 100 kPa was applied to the top surface of the substrate over the radial range of *x* = 0–1 mm, causing upward deformation of the substrate, as shown in Fig. S6A.

The contact interaction between the rigid square and the substrate was defined as frictionless hard contact. The substrate was discretized using four-node bilinear axisymmetric hybrid elements (CAX4H). A global mesh size of 0.1 mm was used, and the mesh was locally refined to 0.01 mm in the contact region and near the axis of symmetry.

After simulation, the deformed top surface profiles (Fig. S6B) were exported to MATLAB (R2024b) for postprocessing. The local surface slope *dh/dx* at a radial position of *x* = 1 mm was calculated by performing a first-order polynomial fit to seven neighbouring points on the deformed profile around the target position. The corresponding slope angle was then obtained as *θ* = tan^*™*1^(*dh/dx*). The computationally obtained angles were compared to those obtained via Eqn. 4 (Fig. S6C).

## 3 Results

### 3.1 Attachment performance on different substrates

We performed 218 force measurements, with an average of 18 measurements per substrate (range: 16–21). Overall, sucker tenacity increased with substrate stiffness and decreased with substrate roughness (Figs. 5, 6A; Fig. S5; Table S2). Relative to the smooth/stiff substrate (model intercept), tenacity on the fine/stiff substrate was not significantly different (estimate = 1.04 ± 0.78, *p* = 0.183), while tenacity on the semi-coarse/stiff and coarse/stiff substrates was significantly lower (estimate = ™1.66 ± 0.84, *p* = 0.050; estimate = ™4.27 ± 0.86, *p* < 0.001, respectively). Substrate stiffness also had a strong effect: suckers on the smooth/soft substrate had significantly lower tenacity than on the smooth/stiff substrate (estimate = ™5.40 ± 0.86, *p* < 0.001), while tenacity on the smooth/medium substrate did not differ significantly from smooth/stiff (estimate = 0.26 ± 0.83, *p* = 0.752). Sucker area was negatively correlated with tenacity across all conditions (estimate = ™0.98±0.27, *p* < 0.001), consistent with smaller suckers generating higher tenacities [19].

**Figure 5:**
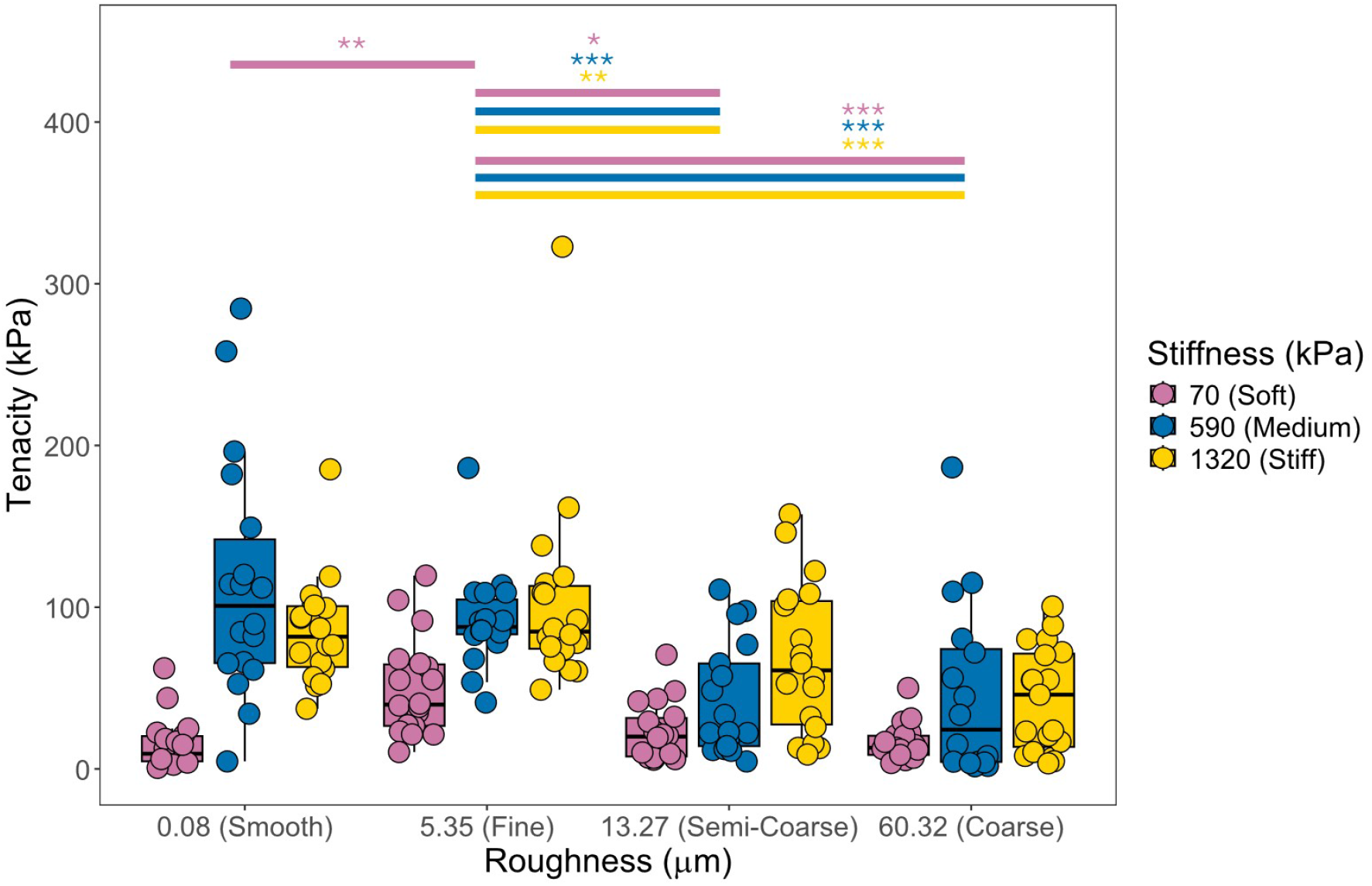
Tenacity *P* of suckers on a range of different substrates differing in stiffness *E* and roughness *Sq*. Data points represent individual measurements, and box plots represent median, quartiles, and 1.5XIQR. Results are color-coded with substrate stiffness (see legend on the right). Bars with asterisks highlight significant differences. (* : p < 0.05, ** : p < 0.01, and *** : p < 0.001.)

**Figure 6:**
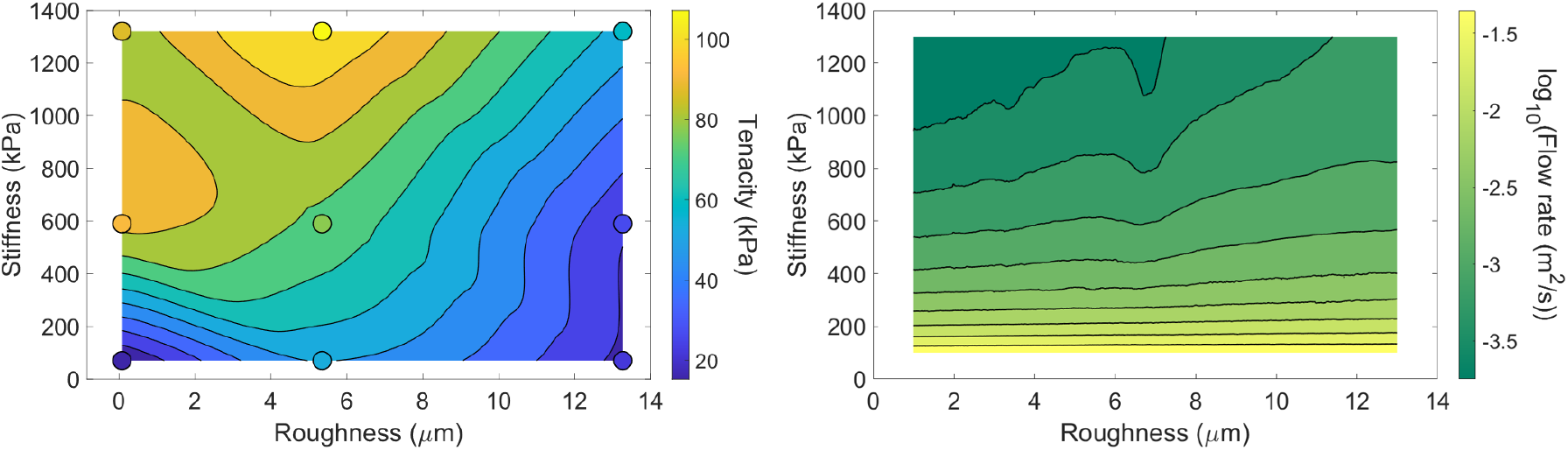
(A) Interpolation of the linear mixed-effects model based on the actual force measurements of tenacity *P* as a function of substrate stiffness *E* and roughness *Sq*. Data points show measurement results, and contour plot show the best fit. (B) Prediction of the flow rate *Q* as a function of substrate stiffness *E* and roughness *Sq* from the mathematical model (Eqn. 5) of leakage through the sucker rim and papillae.

The interaction between stiffness and roughness revealed that the effect of roughness on tenacity depended on substrate stiffness (Table 1). For medium stiffness substrates, roughness had little modulating effect: interactions with fine and coarse roughness were non-significant (estimate = ™1.80 ± 1.16, *p* = 0.123; estimate = 0.04 ± 1.18, *p* = 0.974), and only the semi-coarse roughness showed a significant negative interaction (estimate = ™2.75 ± 1.16, *p* = 0.019), indicating that medium stiffness combined with semi-coarse roughness produced lower tenacity than predicted from the main effects alone.

**Table 1:**
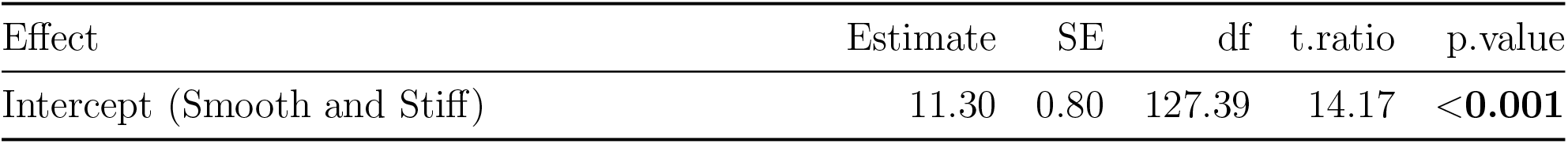
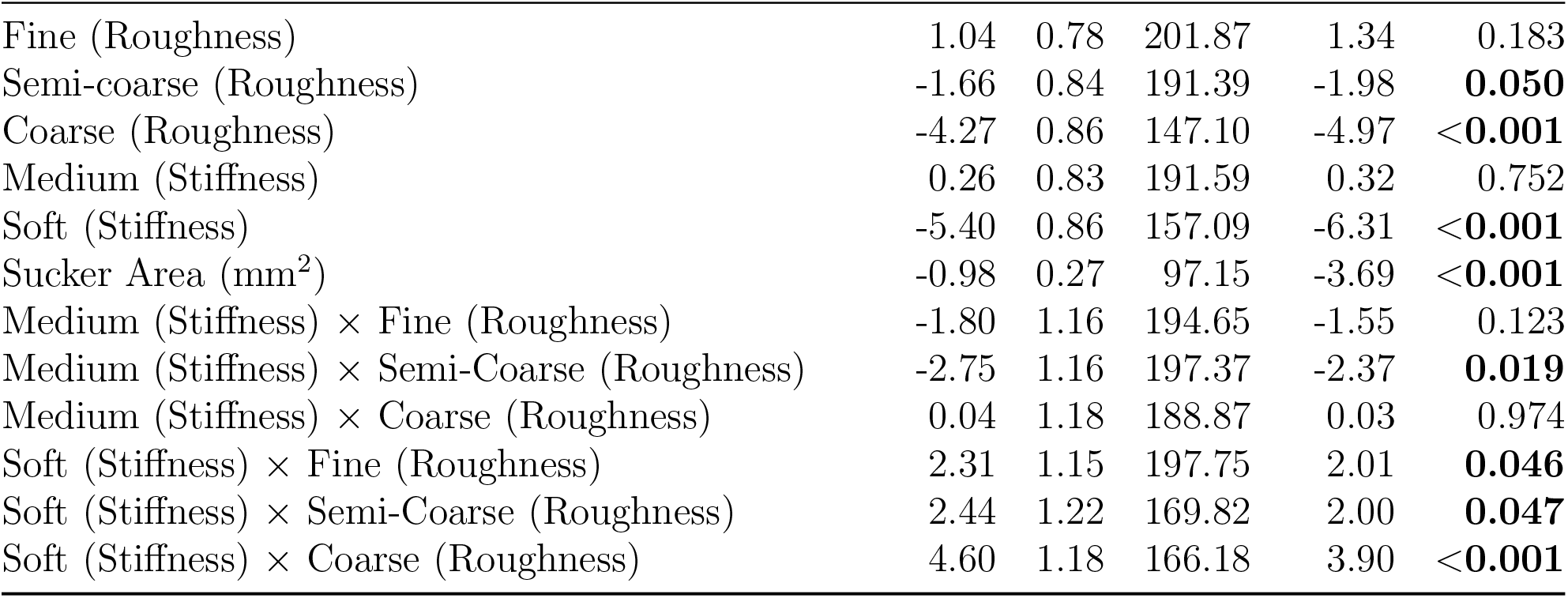
Fixed effects from the linear mixed-effects model examining the influence of stiffness, substrate roughness, and contact area on tenacity (square-root transformed). All estimates represent deviations from the reference condition (smooth/stiff substrate, model intercept). Significant differences are highlighted in bold (*p* < 0.05).

| Effect | Estimate | SE | df | t.ratio | p.value |
| --- | --- | --- | --- | --- | --- |
| Intercept (Smooth and Stiff) | 11.30 | 0.80 | 127.39 | 14.17 | **<0.001** |

| Fixed Effect | Estimate | SE | df | t.ratio | p.value |
| --- | --- | --- | --- | --- | --- |
| Fine (Roughness) | 1.04 | 0.78 | 201.87 | 1.34 | 0.183 |
| Semi-coarse (Roughness) | -1.66 | 0.84 | 191.39 | -1.98 | **0.050** |
| Coarse (Roughness) | -4.27 | 0.86 | 147.10 | -4.97 | **<0.001** |
| Medium (Stiffness) | 0.26 | 0.83 | 191.59 | 0.32 | 0.752 |
| Soft (Stiffness) | -5.40 | 0.86 | 157.09 | -6.31 | **<0.001** |
| Sucker Area (mm<sup>2</sup>) | -0.98 | 0.27 | 97.15 | -3.69 | **<0.001** |
| Medium (Stiffness) × Fine (Roughness) | -1.80 | 1.16 | 194.65 | -1.55 | 0.123 |
| Medium (Stiffness) × Semi-Coarse (Roughness) | -2.75 | 1.16 | 197.37 | -2.37 | **0.019** |
| Medium (Stiffness) × Coarse (Roughness) | 0.04 | 1.18 | 188.87 | 0.03 | 0.974 |
| Soft (Stiffness) × Fine (Roughness) | 2.31 | 1.15 | 197.75 | 2.01 | **0.046** |
| Soft (Stiffness) × Semi-Coarse (Roughness) | 2.44 | 1.22 | 169.82 | 2.00 | **0.047** |
| Soft (Stiffness) × Coarse (Roughness) | 4.60 | 1.18 | 166.18 | 3.90 | **<0.001** |

In contrast, soft substrates showed significant positive interactions across all roughness levels (fine: estimate = 2.31 ± 1.15, *p* = 0.046; semi-coarse: estimate = 2.44 ± 1.22, *p* = 0.047; coarse: estimate = 4.60 ± 1.18, *p* < 0.001). These positive interactions indicate that the reduction in tenacity associated with soft stiffness was partially offset by increasing roughness, particularly on coarse substrates. In other words, roughness had a greater beneficial effect on tenacity for soft substrates than for stiff substrates.

### 3.2 Effects of roughness (given stiffness)

Sucker tenacity decreased on rougher substrates (Fig. 5; Table S1). However, for the soft substrate, sucker tenacity was higher on a fine roughness than the smooth, semi-coarse, or coarse substrates (estimate = 3.35 ± 0.87, *p* = 0.001; estimate = 2.57 ± 0.84, *p* = 0.014; estimate = 3.01±0.73, *p* < 0.001, respectively; Estimated Marginal Means (EMM) Tukey-adjusted). For the medium and stiff substrates, sucker tenacity on the fine substrate was significantly higher than the semi-coarse (estimate = 3.65 ± 0.84, *p* < 0.001; estimate = 2.70 ± 0.84, *p* = 0.008; EMM Tukey-adjusted) and coarse substrate (estimate = 3.46 ± 0.81, *p* < 0.001; estimate = 5.30 ± 0.87, *p* < 0.001; EMM Tukey-adjusted), but not the smooth substrate (estimate = ™0.77 ± 0.82, *p* = 0.789; estimate = 1.04 ± 0.78, *p* = 0.546; EMM Tukey-adjusted).

### 3.3 Effects of stiffness (given roughness)

Sucker tenacity was higher on stiff substrates than soft substrates, for the smooth, fine, and semi-coarse roughnesses (estimate = 5.40 ± 0.87, *p* < 0.001; estimate = 3.09 ± 0.82, *p* < 0.001; estimate = 2.95 ± 0.82, *p* < 0.001; EMM Tukey-adjusted). For the coarse substrate, stiffness did not affect tenacity and sucker performance was in general low (estimate = 0.79 ± 0.87, *p* < 0.64; EMM Tukey-adjusted; Table S2; Fig. S4; Fig. 5). This is also shown in Fig. 6A, where the experimental data is interpolated.

### 3.4 Papillae roughness

Papillae roughness was significantly different from all other substrates except the fine artificial substrate (Fig. 7; *p* < 0.05 for all except for fine, Games-Howell). This is also visualised by cryo-SEM, which shows a close match in roughness at the papillae-fine substrate interface (Fig. 7, Fig. 4C; *p* = 0.569, Games-Howell). Cuttlefish papillae (*Sq* = 6.84 ± 1.83 *µ*m) are marginally rougher than most of the prey species, which were all quite smooth except for the multiple fish scales. Shrimp carapace (*Sq* = 2.00 ± 1.16 *µ*m), the ventral and dorsal crab carapace (*Sq* = 2.14 ± 1.10 *µ*m, *Sq* = 2.11 ± 0.62 *µ*m) and single fish scales (*Sq* = 1.85 ± 0.40 µm) exhibit marginally less roughness than papillae (*p* = 0.08, *p* = 0.035, *p* < 0.001 and *p* < 0.001, Games-Howell, respectively). The roughness of multiple fish scales (*Sq* = 44.80 ± 12.9 *µ*m) was significantly greater than the papillae (*p* < 0.05, Games-Howell).

**Figure 7:**
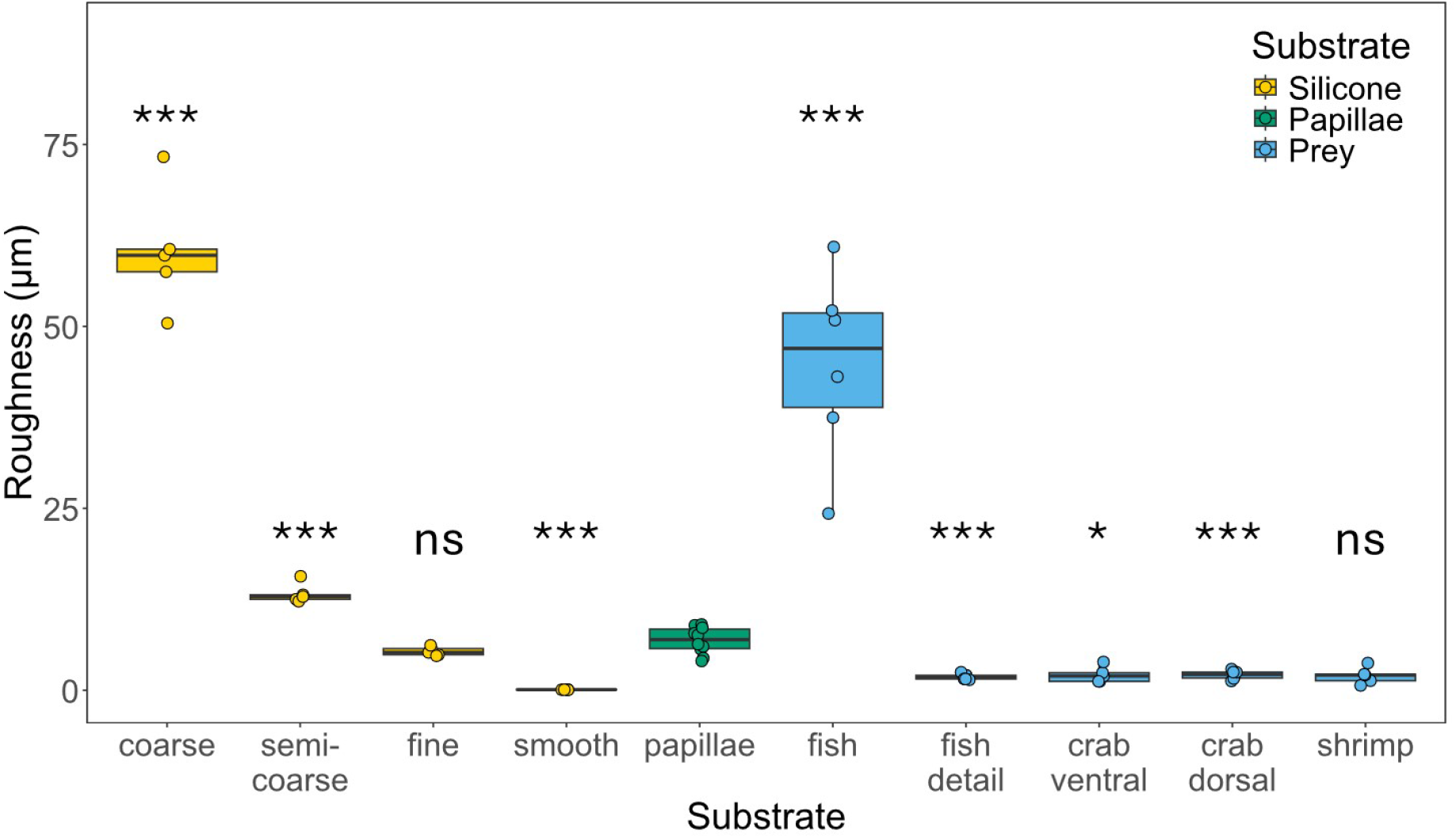
The root mean square roughness *Sq* of the substrates, prey items, and cuttlefish sucker papillae (in yellow, blue and green, respectively; see legend on top right). Sample sizes: Coarse, semi-coarse, fine, smooth and shrimp (n=5); crab ventral, crab dorsal, fish and fish detail (n=6); papillae (n=10). Data points represent individual measurements, and box plots indicate median, quartiles, and 1.5XIQR. A Games-Howell pairwise comparison was performed comparing papillae with all substrates and prey types. Bars with asterisks highlight significant differences. (* : p < 0.05, ** : p < 0.01, and *** : p < 0.001.)

### 3.5 Effect of roughness and stiffness on sucker rim leakage

To examine the interaction at the papillae-substrate interface, we modelled the interlocking of the papillae with substrates of varying roughness and stiffness, assuming pressure-driven plane Poiseuille flow across the sucker rim. The model determined the flow rate of water leakage across the sucker rim. In Fig. 6B, we show how leakage flow rate *Q* varies with substrate roughness *Sq* and stiffness *E*, following Eqn. (5). The model predicts that leakage decreases with increasing stiffness. Softer substrates deform more within the sucker cavity (see Fig. 4A, Fig. S6), increasing both the gap *d* between the papillae and the substrate, leading to a higher leakage flow rate *Q*, and resulting in worse attachment performance. Similarly, a larger roughness increases the gap *d*, thus leading to a higher leakage flow rate *Q*. When papillae and substrate roughness match, the papillae may partially interlock, creating a smaller gap *d* and thus lowering the leakage flow rate *Q*. However, for softer tissues, the substrate deformations increase, diminishing the interlocking effect. The model agrees qualitatively with the interpolated experimental data of attachment performance, or tenacity *P* (Fig. 6A), where lower leakage rate *Q* correlates with higher tenacities *P*.

## Discussion

In this study, we used a combined experimental and modelling approach to study the functional biology of cuttlefish sucker attachment. We measured *ex vivo* cuttlefish arm sucker attachment performance on a range of artificial substrates differing in roughness and stiffness, and used a mathematical model to quantify the microscopic interactions between the sucker rim and the various substrates.

### 4.1 Highest attachment performance when roughness matches

Our pull-off force measurements exhibited the highest tenacities with substrates that had roughnesses that matched the average papillated rim roughness. These roughnesses match common prey, crabs and shrimp [2]. Of the prey items analysed in our study, fish scales exhibited the highest roughness values, which could make attachment more difficult. However, fish scale roughnesses can vary significantly among species, and our tested fish scales are likely not representative of all fish [2].

The trends we observed for attachment performance are congruent with studies of northern clingfish, which examined grain sizes ranging from 0 *µ*m to 2000 ™ 4000 *µ*m. Within this range, Northern clingfish adhered best to substrates with grain sizes between 35 ™ 269 *µ*m—-similar to their average papillae size (∼ 130 *µ*m). Moreover, bioinspired micro-structured suction cups mimicking the papillae performed best on roughnesses that match the size of their artificial papillae structures [11, 23].

Our mathematical model of leakage through the sucker rim provides a mechanism that might explain these findings. The stronger attachment onto the substrate with fine roughness (*Sq* = 5.35 *µ*m) may be due to interlocking of the papillae, as previously proposed for clingfish [11, 12]. This promotes intimate contact between substrate and sucker, and helps create a tighter seal around the sucker rim. Furthermore, previous research showed that the closer contact increases viscous forces due to Stefan’s adhesion when mucus is present [11].

Our findings suggest that cuttlefish suckers perform best when the roughness of the substrate matches that of their papillae. It, therefore, seems plausible that cuttlefish papillae may have evolved to match the substrate roughness of their prey. At the scale of the papillae, mechanical interlocking may be essential to resist collapse at high differential suction pressures. Similar observations have been made in clingfish, which exhibited lower maximum suction on smooth substrates than on rough substrates [12]. Likewise, in suckermouth armoured catfish (Loricariidae), the length of the papillae on their sucking mouthparts is hypothesised to be adapted for different substrate types like sand, wood, and/or stone [13]. Future work could examine other cephalopod species with other prey preferences to see if their roughnesses match their specific prey.

### 4.2 Higher attachment performance on stiffer substrates

Our experimental force measurements showed that sucker tenacity increased with increasing substrate stiffness. Given that crabs, shrimp, and fish scales are very stiff, with elastic moduli on the order of *E* ∼ GPa [7, 36, 37], we expect that cuttlefish can adhere quite well to these species. On the other hand, other cephalopod species are much softer (*E* = 500 ™ 1200 kPa for the muscular mantle of squid *Loligo pealei* [38]). Therefore, we expect roughly the same performance as we measured on the medium and stiff substrates. From JKR contact theory of a more simple geometry, i.e., a flat, cylindrical punch in contact with an elastic substrate, the adhesive force is expected to increase with substrate stiffness, or ∼ *E*^1*/*2^ [16]. However, cuttlefish sucker contact morphology is more complex and the presence of high suction pressures (∼ 100 kPa) leads to a more nuanced situation.

Our mathematical model of leakage through the sucker rim addresses the complex morphology and provides insight into the potential underlying contact mechanics. The model shows that weaker attachment onto soft substrates can be explained by deformation as a result of low pressures within the sucker cavity. That is, as the pressure decreases inside the sucker, the substrate is sucked into the inner volume of the sucker. Computational finite element modelling further confirms that soft substrates get pulled into the sucker cavity due to the suction pressures (Fig. S6A-B). Such deformations may promote separation between the sucker rim and substrate.

Additionally, soft substrates are less effective in expelling water from the contact interface than stiff materials, resulting in a less effective seal. Previous observations on the contact mechanics of soft substrates underwater found that pockets of water can be trapped at the interface between two deformable substrates, thus hindering adhesion and friction [9].

### 4.3 Limitations and scope

#### 4.3.1 Mathematical model

While our mathematical model can largely predict sucker performance, it cannot predict the significant increase in sucker performance for soft substrates with a fine roughness. In our mathematical model, soft substrates deform under a pressure differential, which, in addition to a larger average gap size, also leads to the substrate sliding into the sucker cavity. This sliding at the interface could introduce a lubricating layer of water beneath the sucker rim. This is also observed in clingfish that adhere worse on fouled substrates, because the lower friction is thought to lead to inward slippage of the sucker [39]. We hypothesise that, in the case of a matching roughness in soft substrates, the close contact of papillae and the fine substrate results in higher friction and, thus, less sliding along the substrate and higher suction performance.

Our simplified mathematical model neglects the local deformations of asperities on the substrates. Previous models have predicted that elastic asperities would flatten out during contact [40], which may increase substrate conformability and decrease leakage through the sucker rim. Therefore, we expect that our model overestimates leakage on softer substrates, but, qualitatively, the model still captures the interlocking of papillae and substrate asperities.

In our experiments, we measured tenacities as large as 300 kPa, but, in our model, we assumed a suction pressure of 100 kPa. The tenacities in the experiments were calculated using the inner diameter of the sucker cavity aperture. This assumes that pressure gradient from inside to outside the sucker occurs instantly at the inside edge of the sucker rim. This may result in an underestimation of the sucker area, and thus an overestimation of tenacities (and inferred suction pressures). Since the larger tenacities were observed for only a small subset of the trials with the stiffer substrates, this assumption in our model would underestimate the leakage rate for the stiffer substrates. However, we would expect a similar qualitative trend in leakage rate.

Finally, the deformation of the soft substrates into the sucker cavity was assumed to be linear, leading to a simplified equation for the tilt angle of the substrate, Eqn. (4). Using a computational finite element model of a linearly elastic substrate experiencing suction pressure, we found a similar relationship between tilt angle *θ* and substrate stiffness *E* (Fig. S6C). Therefore, our simplified representation still captures the complex deformations caused by local suction pressures. Both methods for estimating substrate tilt angle likely overestimate these values, as we expect the substrate to be attracted to the papillae. However, the exact nature of these interactions are unknown, and could be due to suction pressures, viscous forces due to Stefan’s adhesion when mucus is present [11], or potential van der Waals interactions.

#### 4.3.2 Experiments

Although our experiments show trends for roughness and stiffness, our study was limited to passive sucker attachment on animals that have been frozen. The role of active muscles during attachment remains unclear and may be critical for substrates that are difficult to adhere to. In our experiments, muscles do play a passive role, as their elasticity matters for sucker performance. However, our measurements show that the performance of previously frozen suckers is comparable to fresh suckers (with reflexive muscle activity; see Fig. S1B). Still, the forces generated by the suckers of living cuttlefish on varied substrates remain unknown and our work here represents the first attempt at understanding this complex system.

Additionally, our work only examined pull-off force in the normal vector. We observed asymmetry in both the morphology of the suckers as well as the papillae, which suggests that they are adapted to function directionally (Fig. 3E). This asymmetry, and its functional role in generating friction or resisting shear forces, has been documented in the spinules of remoras and the papillae of northern clingfish, and has also been experimentally evaluated in artificial bionic papillae [11, 21, 41]. Future work could explore cuttlefish attachment *in vivo* and test how this asymmetric directionality affects friction.

The higher attachment performance on soft substrates with roughness that matches cuttlefish papillae provides an interesting avenue for bioinspired suction cups. Previous research has focused successfully on how to attach to rough substrates, using compliant suction cups [14, 23, 42]. However, strong attachment to soft substrates is still difficult, let alone in combination with a rough exterior. Our results suggest that a possible biomimetic solution is to use stiff artificial papillae that match the roughness of the substrate. This agrees with previous research that suggests that sucker attachment performance increases with increasing effective elastic modulus (i.e., the combination of the elastic moduli of the substrate and sucker) [14]. The incorporation of stiff papillae may improve the versatility of bioinspired suction cups, especially when functioning underwater.

In summary, this work suggests that papillae may enhance attachment performance by reducing the effective volume between the sucker rim and the substrate. This conclusion is supported qualitatively by a mathematical model that represents papillae and substrate rugosity as variable sine waves and incorporates deformations due to suction pressure. Taken together, these results suggest that the papillated rims of cuttlefish suckers may be adaptively tuned for adhering to prey with comparable substrate roughness. More broadly, this work provides new insight into the functional adaptations of cephalopod suckers and offers a foundation for the design of future biomimetic suction cups.

## Acknowledgements

We thank Julia Jansen Holleboom and Henk Schipper for the histological images, Hendrik du Toit for providing the sole (*Solea solea*) samples, Alexander Koehnsen and Johan van Leeuwen for feedback on early drafts and fruitful discussions, Lia Hemerik and Johannes Ransijn for support on the statistical analysis, Remco Pieters and Anne de Waal for technical support for the pull-off force experiments.

## Author Contributions

Conceptualization: BKvO, GJA, JStL; Methodology: BKvO, GJA, JStL, TG, EtLB, MG; Formal analysis: BKvO, GJA, JStL, EtLB, MG, BZ, FTM, SWSG, TG; Investigation: BKvO, GJA, JStL, TG, EtLB, MG, BZ, YL; Resources: GJA; Writing – original draft: BKvO, JStL; Writing – review & editing: BKvO, GJA, JStL, TG, EtLB, SWSG, FTM; Visualization: BKvO, GJA, JStL, EtLB, MG; Supervision: BKvO, GJA, SWSG, FTM; Project administration: GJA; Funding acquisition: BKvO, GJA.

## Data Availability

Data and analysis are available at Figshare: https://figshare.com/s/e15b0dc2e74f3c8e28d8

